# CRISPR-Cas12a-Based Rapid and Sensitive Detection of *rpoB*L378R in *Mycobacterium tuberculosis*

**DOI:** 10.1101/2023.06.06.543922

**Authors:** Li Yang, Xiaoyu Li, Jing Tang, Yue Zhu, Kai Ma, Yuma Yang, Zhaoyuan Hui, Yanyan Qin, Hetian Lei, Minghai Shan, Yanhui Yang

**Author notes:** Correspondence Hetian Lei: 18 Zetian Road, Fudian District, Shenzhen, China.; Minghai Shan: The General Hospital of Ningxia Medical University, Yinchuan, China; Yanhui Yang: The School of Basic Medical Sciences, Ningxia Medical University, Yinchuan, China.

## Abstract

Rifampin is the most effective drug in the treatment of tuberculosis, whose major pathogen is *Mycobacterium tuberculosis* (MTB), whereas there are still certain MTB strains resistant to the therapy of rifampin. The *rpoB* mutations play a central role in MTB resistance to the rifampin therapy, so it is crucial to identify these mutations in order to discover novel therapeutic approaches to these drug-resistant MTB strains. Here we show that a CRISPR-Cas12a-based detection platform with recombinase polymerase amplification and fluorescence reporter can be utilized to detect and visualize an MTB drug-resistant point mutation (*rpoB*<u>L378R</u>) from its *rpoB* wild type. Notably, this detection system is highly specific because it did not cross-react with contrived reference samples containing the genomes of MTB *H37Rv*, *Mycobacterium smegmatis* (*M. smegmatis*), *Mycobacterium aureus* (*M. aureus*), and *Escherichia coli* (*E. coli*). Collectively, this strategy based on CRISPR-Cas12a that we show in this report is simple, sensitive as well as specific for detection of the rifampin-resistant MTB *H37Rv* with the *rpoB*L378R mutation, indicating that this CRISPR-Cas12a-based detection platform has high potential to be exploited for clinic application to identify MTB mutations.

## Importance

The CRISPR-Cas12a-based platform is simple, sensitive as well as specific for detection of the rifampin-resistant MTB *H37Rv* with the *rpoB*L378R mutation. CRISPR-Cas12a system combined with recombinase polymerase amplification to improve detection sensitivity, and avoid using of PCR devices to simplify the application of expensive equipment. Fluorescence readout and lateral flow detection were used to achieve rapid, portable, wide compatibility, highly sensitivity and highly specificity in CRISPR-Cas12a system.

## Introduction

Tuberculosis (TB) that is caused by the infection of *Mycobacterium tuberculosis* is one of the major infectious diseases globally[1]. According to the WHO Global Tuberculosis Report 2021, there are 9.9 million TB patients and 1.5 million people died from this disease worldwide[2]. While 85% of patients with TB can be cured after drug treatment for 6 months, the rest carry drug-resistant TB (DR-TB). The emergence of DR-TB, especially multidrug-resistant TB (MDR-TB), poses an enormous challenge to the prevention and treatment of TB[2]. The development of drug resistance is a direct result of gene mutations that need to be identified by PCR or sequencing technology[3]. Many first-line clinical drugs place selective pressure on bacteria to evolve drug-resistance at various genotypes conferring drug-resistance, such as rifampicin (RIF) in *rpoB* (*e.g.*, codons S450L, L378R, L430P, H445N), and isoniazid (INH) in genes *katG* (codon S315T) [4-6]. *rpoB* is the target gene of Xpert MTB/RIF detection. The mutations of S450L and H445R/Y/L in RIF resistance determination region and L378R and R871H outside RIF determination region are mainly single base mutations related to resistance of these first-line TB treatment drugs. To reduce the complexity and cost of drug resistance diagnosis to a certain extent, it is crucial to develop a rapid, sensitive and accurate TB detection approach for the prevention and treatment of TB, especially MDR-TB.

Advances in nucleic acid detection techniques have contributed to the prevention and treatment of drug-resistant tuberculosis. The advantages of accurate and sensitive detection have led the world health organization (WHO) extensively acknowledge nucleic acid detection technologies, which have replaced previous microscopic technology as an important TB testing standard. Polymerase chain reaction (PCR) is the most widely used for pathogen nucleic acid detection as PCR is higher accuracy and specificity[7-9]. However, there are certain drawbacks in using PCR including the requirement for expensive equipment and experienced technicians. Additional methods have also been reported for nucleic acid detection including Xpert MTB/RIF, DNA sequencing, and line probe assay (LPA)[10, 11]. Nevertheless, the disadvantages of expensive, instruments-dependent, or time-consuming restrict them in economically backward areas or outdoor environments. Recently, several isothermal amplification methods, such as RPA[12, 13] and loop-mediated isothermal amplification (LAMP)[14], have been developed and expected to reduce the restriction of PCR instrument-dependent amplification. Although RPA and LAMP have been successfully developed for detection of ASFV and HBV [13, 14], their detection accuracy depends on specificity between the primers and templates[15].

Nucleic acid detection can also be accomplished with CRISPR-Cas systems, in which many Cas proteins including Cas12a, Cas12b, Cas13a and Cas14 are able to recognize dsDNA, ssDNA or RNA sequence under the guidance of single guide (sg) RNA [16-20]. CRISPR-Cas nucleic acid detection technology has been successfully applied for examining many diseases including Hepatitis B Virus (HBV)[21], SARS-CoV-2[17, 22], HIV[17], Dengue and ZIKA[23, 24]; in addition, CRISPR-Cas-based detection technologies have also achieved single nucleotide recognition [24-26]. Therefore, CRISPR-Cas12a in combination of recombinase polymerase amplification (RPA) can be utilized to establish simple, fast, highly sensitive and specific detection methods for detecting the rifampin-resistant MTB *H37Rv* strains with the mutations such as *rpoB*L378R [27, 28].

In this study we hypothesized that the MTB drug resistance mutant gene *rpoB*L378R could be used as a detection target, and the CRISPR-Cas12a system could be used to detect single base differences in the mutant gene sequence. Our aim of this investigation was to establish a rapid, convenient, economical, highly specific, highly sensitive and visual detection method for MTB drug resistance point mutation by combining RPA isothermal amplification technology, fluorescence detection and paper chromatography, and layout a foundation for further development of MTB drug resistance mutation detection reagents in clinic application.

## 1 Materials and Methods

### 1.1 Materials

Major reagents involved in this study included EnGen LbCas12a (Cpf1) (New England BioLabs, M0653T), plasmid mini kit I (OMEGA, D6943-02), Luria broth (LB) media, Ampicillin (100 mg/mL), DH5α competent cell, NEBuffer2.1, polyacrylamide gel electrophoresis Gel Fast Preparation Kit (EpiZyme, PG114), Gel Extraction Kit D2500 (OMEGA, D2500-02), TaKaRa MutanBEST Kit (Takara, R401), QuickCutTM Hind III (TAKARA, 1615), TwistAmp^®^ Basic Kit (TwistDx, TABAS03KIT), Milenia HybriDtect1 (TwistDX, Milenia01), UltraPure DNase/RNase-Free Distilled Water (Thermo, 10977015) and RNase inhibitor (Takara, 2313A). Thermo Fisher Nanodrop 1000 Spectrophotom was used to quantify nucleic acid. Fluorescence signals were detected by PerkinElmer EnVision Multimode Plate Reader, and fluorescence imaging was gained by Bio-Rad GEL Doc XR.

### 1.2 Methods

#### 1.2.1 Target sequence and crRNA preparation

We selected *rpoB* gene sequence (from 760794 to 761540, long 753 bp) (GenBank accession NC_000962) as a research object. The wild-type *rpoB* gene of MTB *H37Rv* was downloaded from National Center for Biotechnology Information, and subsequently it was synthesized and cloned into topo plasmid (topo-*rpoB*) by Beijing Ruibiotech. Both ends of *rpoB* sequence were synthesized with restriction enzyme cutting sites for *Hind* III. To obtain *rpoB*L378R (CTG>CGG) target sequence, the topo-*rpoB* plasmid was performed a point mutation in codon 378 with mutation primers (*rpoB*L378R_mut_F and *rpoB*L378R_mut_R) and TaKaRa MutanBEST Kit’s introduction. All of sequences were exhibited in Table 1.

**Table 1.**
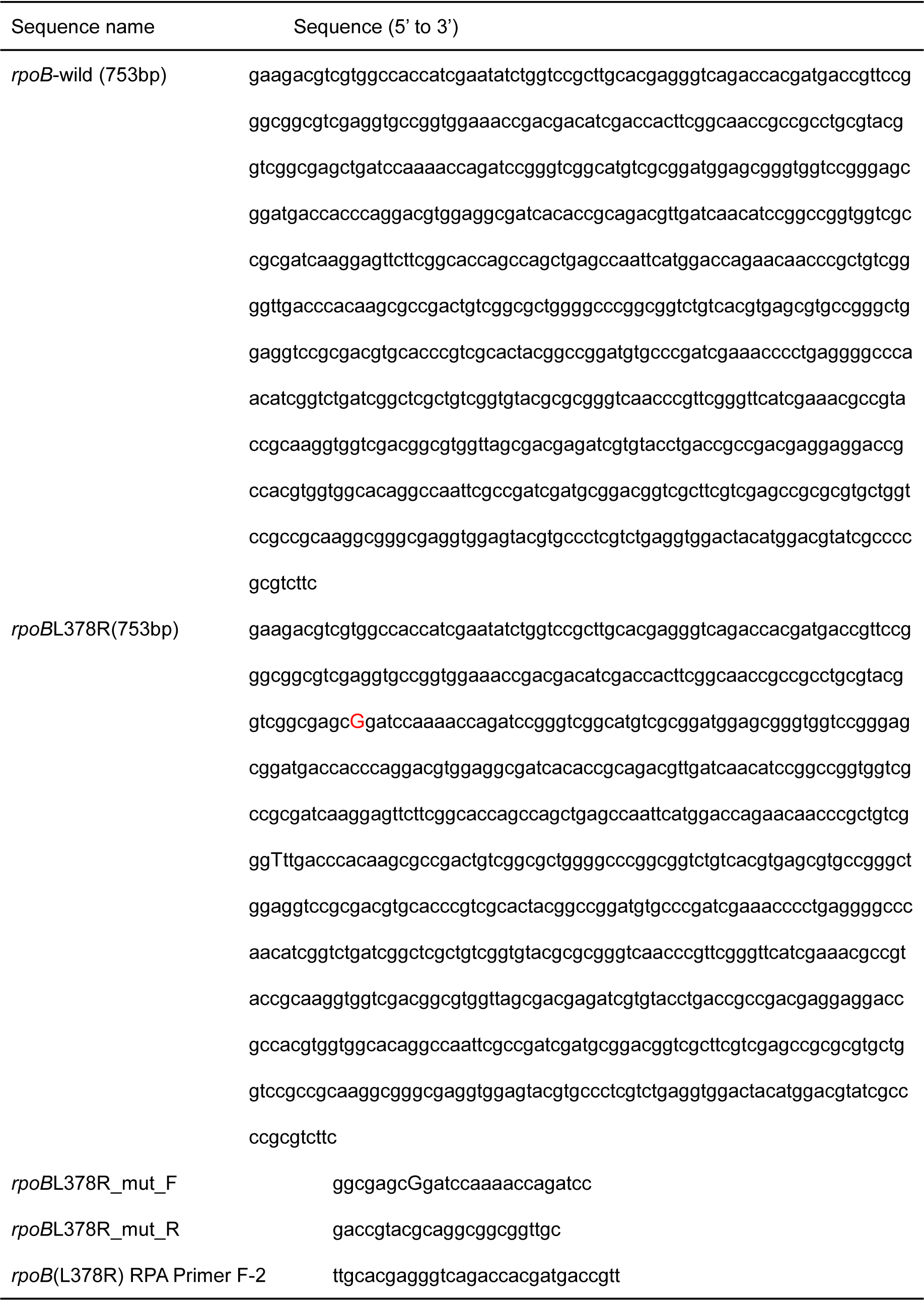
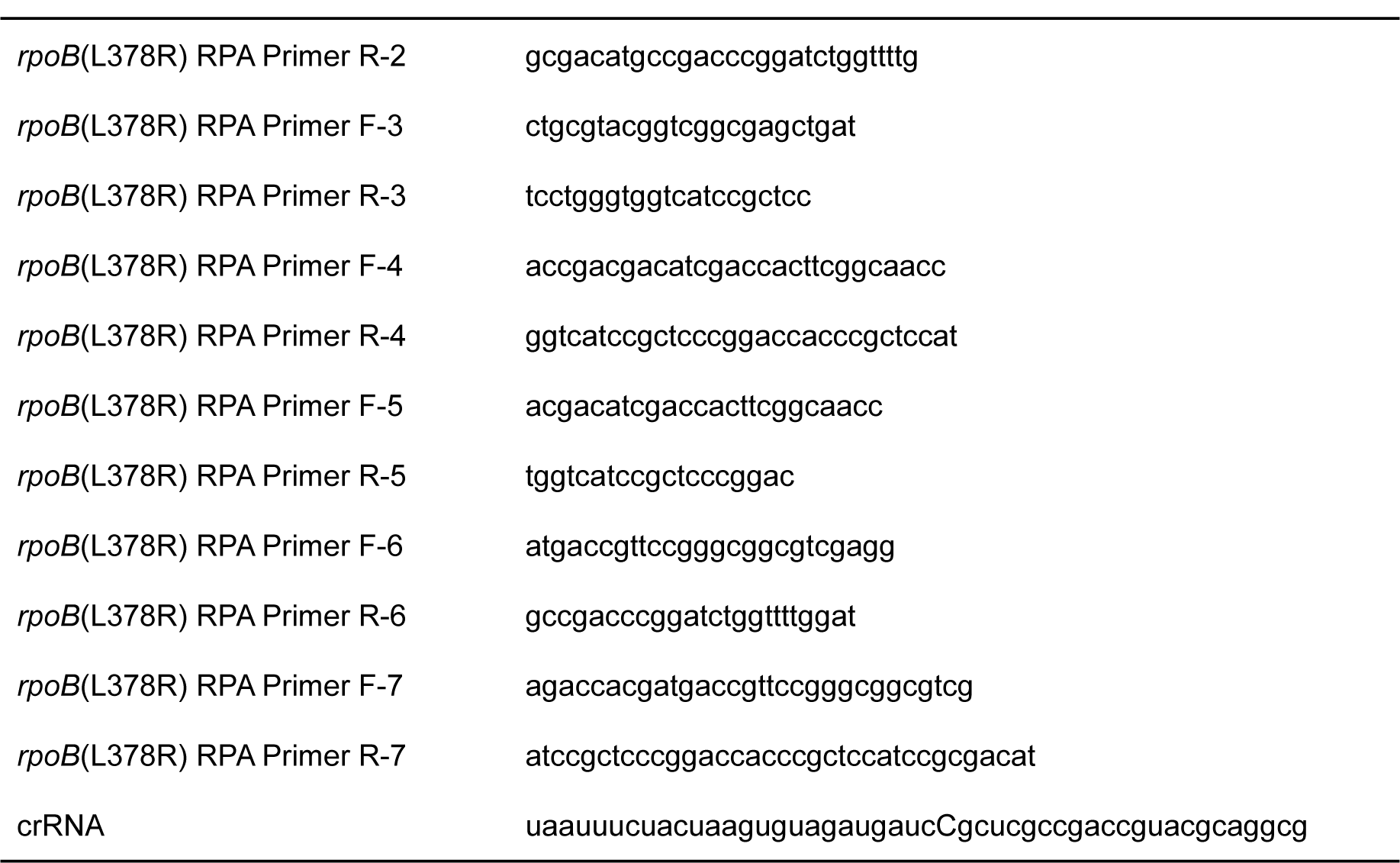
The sequence list of primers, crRNAs and *rpoB*

The crRNA was designed to pair with *rpoB*L378R mutant sequence by CRISPOR website (http://crispor.tefor.net/crispor.py), and the principle of designing crRNA was described previously[27]. Above-mentioned all nucleic acid sequences were synthesized from Beijing Ruibiotech.

#### 1.2.2 Design of RPA amplification assay and primers

To improve LbCas12a detection efficiency and sensitivity, RPA was performed with TwistAmp® Basic Kit. Four RPA primer pairs were designed at both ends of a mutant site of *rpoB*L378R target sequence used Primer Premier 5.0 and refered to the TwistAmp Assay Design Manual. RPA primers length was designed from 19 nt to 35 nt, and the producing amplicon size was under 150 bp. Then, RPA primers were confirmed further by NCBI Primer-BLAST based on the RPA primers design principles. Before RPA assay, we selected the best primer pair based on the ratio and specificity of amplification, and then RPA amplification was performed strictly according to TwistAmp^®^ Basic Kit’s introduction. The primer sequences were synthesized from Beijing Ruibiotech, and listed in Table 1.

#### 1.2.3 LbCas12a cleavage assay

LbCas12a cleavage assay was carried out based on a previous protocol [25] with slight modifications. Briefly, Cas12a cleavage reaction was implemented with 400 nM Cas12a, 400 nM *rpoB*L378R-targeted crRNA, 1U/μL murine RNase inhibitor, and 100 ng *rpoB*L378R or *rpoB*_wild type in 1 NEB buffer 2.1. This mixture was incubated at 37℃ for 60 min, and then analyzed by 2% agarose gel electrophoresis.

#### 1.2.4 FAM-BHQ1 fluorescence reporter detection

The ssDNA FAM-BHQ1 reporters contained a poly (T) oligonucleotide sequence linked to fluorophore and quencher. In the FAM-BHQ1 fluorescence reporter detection assay, LbCas12a firstly bonded crRNA to form a LbCas12a-crRNA complex, and LbCas12a was activated when crRNA paired with the target DNA sequence, triggering the trans-cleavage activity of LbCas12a to cleave a non-target ssDNA reporter. In this assay there were 400 nM LbCas12a, 400 nM crRNA, 1 U/μL murine RNase inhibitor, 2.4 μM ssDNA FAM-BHQ1 fluorescent reporter and 100 ng target DNA *rpoB*L378R or *rpoB*_wild type in 1 NEB buffer 2.1. The reaction mixtures were quickly transferred to a light-proof 96-well plate at Multimode Plate Reader preheated 37 ℃, and fluorescence signals were measured every 5 min (FAM: λ_ex_=492, λ_em_=517). In addition, we also observed fluorescence intensity under light including UV, LED blue and no excitation light. In optimization of fluorescence detection conditions, we tried to improve fluorescent detection condition including incubating time (every 10 min for a total of 60 min), concentration of RPA primers (0.24, 0.36, 0.48, 0.60 μM) and ssDNA FAM-BHQ1 (1.6, 3.2, 4.8, 6.4 μM), and proportion of crRNA and Cas12a (1:1, 1:2, 2:1 based on 400 nM) to maximize detection effect.

#### 1.2.5 Lateral flow readout of Cas12a detection

In lateral flow detection assay, the reaction mixture contained 400 nM LbCas12a, 400 nM crRNA, 1U/μL murine RNase inhibitor, 1 μM ssDNA FAM-biotin reporters,RPA reaction mix of *rpoB*L378R or *rpoB*_wild and 1×NEB buffer 2.1type. After incubation for 40 min, 100 μL of HybriDetect 1 assay buffer (from Milenia HybriDetect 1 kit) was added. Subsequently, lateral flow detection was performed by placing a HybriDetect 1 lateral flow strip into a reaction system. After 1-2 min the colored readout was developed.

#### 1.2.6 Preparation and analysis of the bacterial samples

*rpoB*-negative MTB *H37Rv* strains were inactivated at 100 ℃ for 10 min, and the genome of *M. Smegmatis*, *M. aureus*, *E. coli* and MTB *H37Rv* were extracted using Bacterial DNA Kit and preserved at −80 ℃. The *M. Smegmatis*, *M. aureus* and *E. coli* genomes were mixed with *rpoB* wild-type or *rpoB*L378R fragments at a molar ratio of 1:1, respectively. The content of each genome in the mixture system was 1 nM. Then, fluorescence reporter detection was performed, and there were 400 nM LbCas12a, 800 nM crRNA, 1 U/μL murine RNase inhibitor, 4.8 μM ssDNA FAM-BHQ1 fluorescent reporter and 1 nM mixture of *rpoB*L378R or *rpoB*_wild type and the genome in 1 NEB buffer 2.1, see 1.2.4 for details. And lateral flow detection assay, there were 400 nM LbCas12a, 800 nM crRNA, 1 U/μL murine RNase inhibitor, 1 μM ssDNA FAM-biotin reporters and 1 nM mixture of *rpoB*L378R or *rpoB*_wild type and the genome in 1 NEB buffer 2.1, and see 1.2.5 for details. Furthermore, we added MTB *H37Rv* strains and HEK293T genome to further confirm the assay.

#### 1.2.7 Statistical Analysis

Representative data were the mean ± standard deviation (SD) from triplicates. Statistical analyses were performed using GraphPad Prism 8.0.2 using two-way ANOVA. Significance was considered at \**p* < 0.05, \*\**p* < 0.01, \*\*\**p* < 0.001, \*\*\*\**p* < 0.0001; ns indicated no significance.

## 2 Results

CRISPR-Cas12a can be leveraged to detect a single nucleic acid mutation. Our goal of this research was to establish a rapid, convenient, economical, highly specific, highly sensitive and visual method for detecting MTB drug resistance point mutation. Thereby, we hypothesized that the CRISPR-Cas12a system could be employed to detect the MTB drug resistance mutant genes including *rpoB*L378R.

### 2.1 dsDNA cleavage of *ropB*L378R based on CRISPR-Cas12a

To accomplish the CRISPR-Cas12a cleavage detection on dsDNA *rpoB*L378R as shown in Figure 1A, LbCas12a and crRNA were mixed together to form a LbCas12a-crRNA complex in 37°C, the *cis*- and *trans*-acting cleavage activity of LbCas12a that could achieve targeted and no-targeted cleavage in turn was activated when crRNA and *rpoB*L378R bases paired perfectly[25]. Here, *rpoB*L378 was mutated to L378R (CTG>CGG) (Figure 1B). To test the *cis*-cleavage activity of LbCas12a for *rpoB*L378R compared to *rpoB*_wild type, mutant specific crRNA was designed to distinguish *rpoB*L378R from *rpoB*_wild type (Figure 1C). *Cis*-acting cleavage lets a *rpoB*L378R sequence (753bp) divide into two sections shown by 2 % agarose gel electrophoresis, resulting in a shorter band only shown in lane 4 (Figure 1C). This shorter DNA fragment represented a positive result from the cleaved *rpoB*L378R in the LbCas12a cleavage assay.

**Figure 1.**
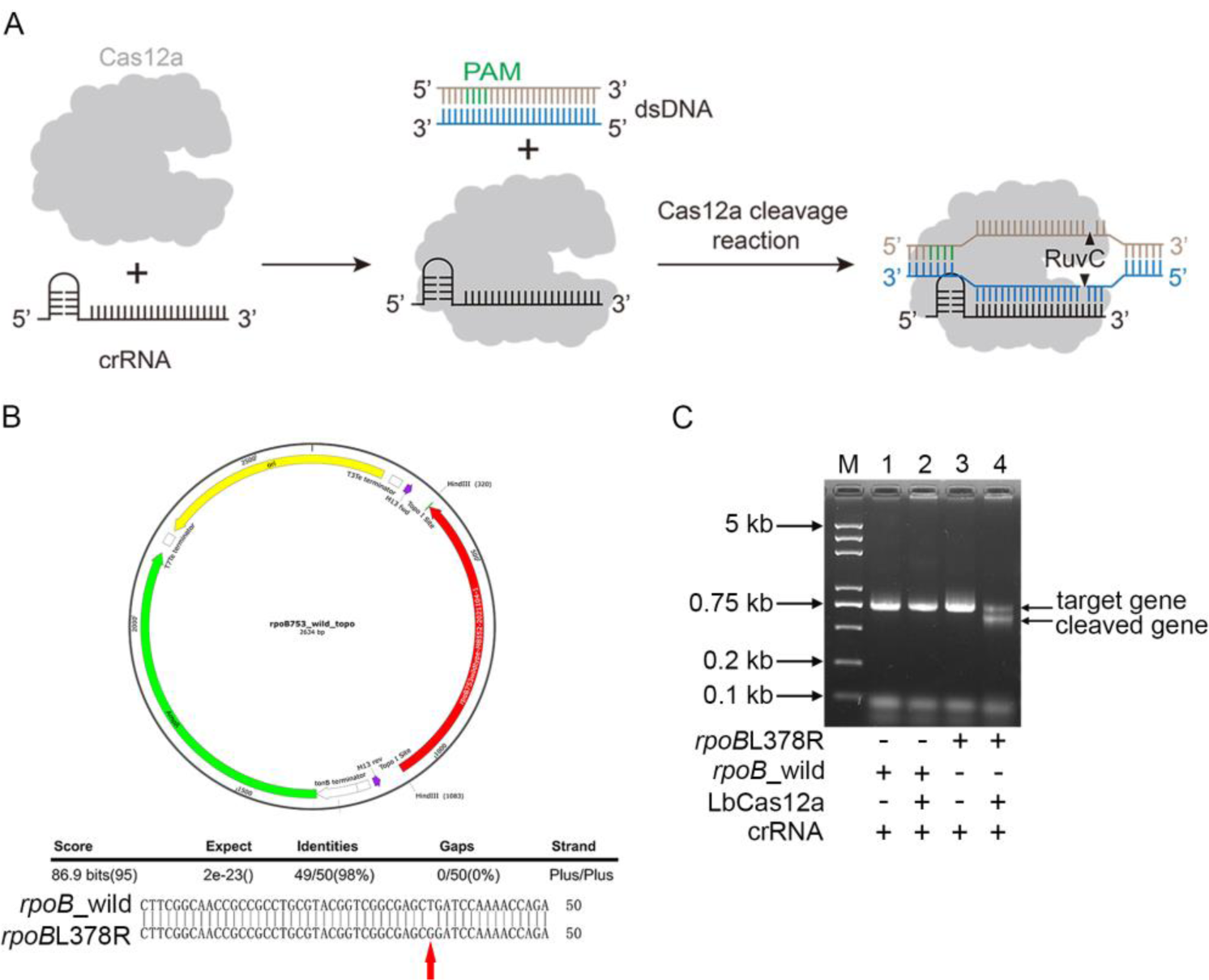
dsDNA Cleavage of crRNA-guided LbCas12a. **A.** Schematic of the components of CRISPR-Cas12a system and its cleavage principle. PAM: Protospacer Adjacent motif, dsDNA: double strand DNA. **B.** The profile of *rpoB* plasmid and sequencing result of *rpoB*L378R point mutation. **C.** Verification of cleavage activity of CRISPR-Cas12a in *rpoB*L378R. M: DL5000 DNA marker, 1-4: *rpoB*_wild, *rpoB*_wild+CRISPR-Cas12a, *rpoB*L378R, *rpoB*L378R+CRISPR-Cas12a. The two cleavage products by LbCas12a under the crRNA guidance are 135 bp and 618 bp.

### 2.2 Detection of *rpoB*L378R with CRISPR-Cas12a is simple, rapid and specific

To visually identify *rpoB*L378R, LbCas12a trans-cleavage activity (collateral activity) was subjected to a non-targeted cleavage ssDNA FAM-BHQ1 fluorescence reporter assay (Figure 2A). In this assay, the RPA primers were firstly optimized. As shown in Figure 2B, there was a brighter and denser single band at about 150 bp in lane 6, indicating this pair of RPA primers resulted in the best specificity and amplification efficiency among the tested pairs. Thereby, this pair of primers was used in the following experiments. We next verified LbCas12a trans-cleavage activity to a non-target DNA sequence (Figure 2A), and tested the specificity of point mutation discrimination by combining a ssDNA fluorescence reporter. As shown in Figure 2C, reaction 3 with *rpoB*L378R sequence, crRNA and LbCas12a produced stronger fluorescence signal than that with *rpoB*_wild in reaction 7, and a fluorescence signal could be directly seen under blue LED or UV light. In addition, the specificity of the generated fluorescence signal was confirmed by a polyacrylamide gel electrophoresis (PAGE) assay as shown in Figure 2C. There was a thick stripe with shorter DNA size in lane 3 (Figure 2C), which represented the cleaved ssDNA FAM-BHQ1 reporters with a strong fluorescence signal. In other reaction conditions there were only weak bands with relatively longer DNA sizes in corresponding lanes, which might be attributed to fluorescence quench of the intact uncut reporter. These results indicate that LbCas12a detection assay provides a simple, rapid and specific detection of *rpoB*L378R.

**Figure 2.**
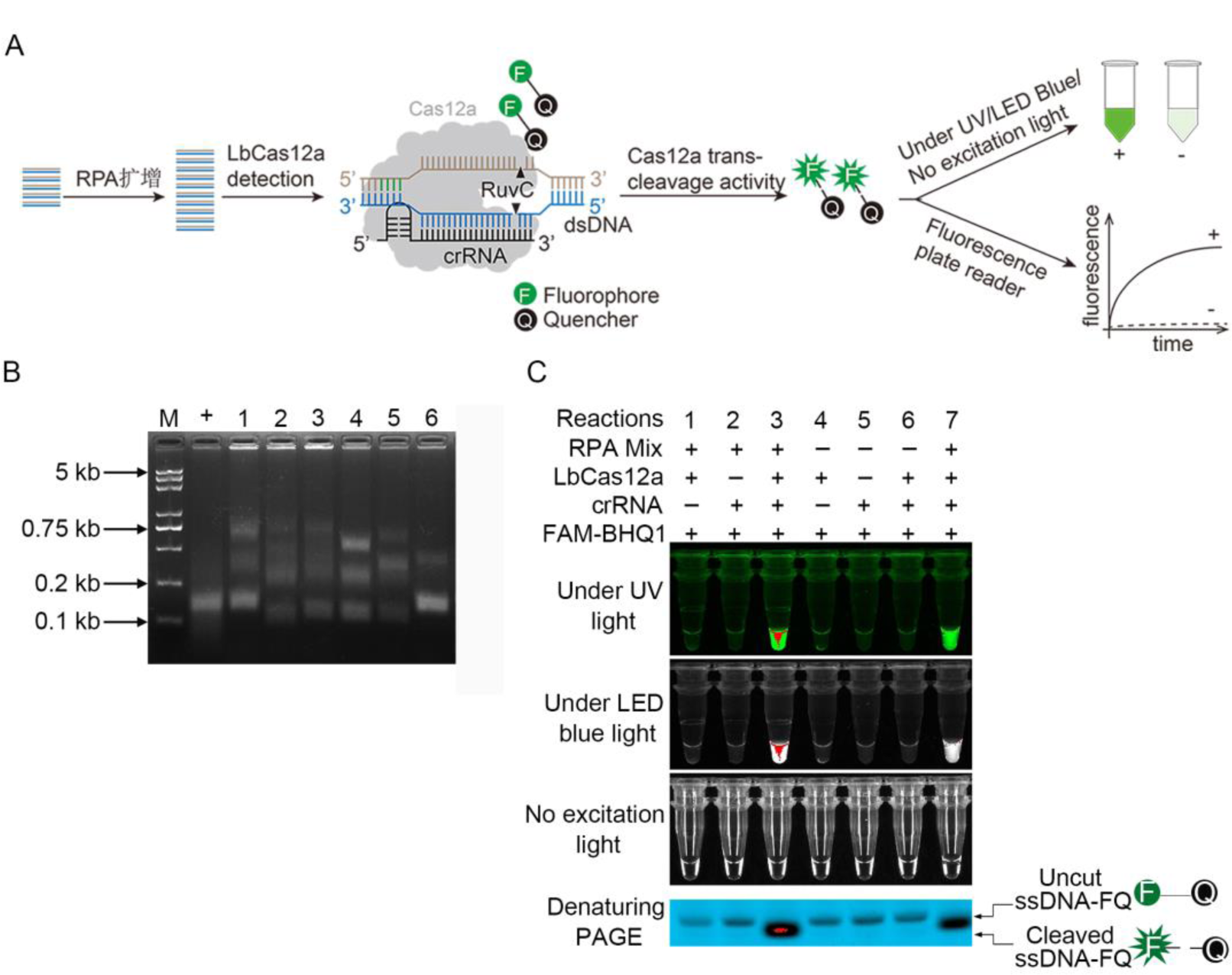
CRISPR-Cas12a-based nucleic acid detection was carried out by fluorescence signal readout. **A.** Schematic of CRISPR-Cas12a-based fluorescence signal readout. RPA: Recombinase Polymerase Amplification, **+**: positive, **-**: negative. **B.** Screening result of RPA amplification primers. M: DL5000 marker, 1-6: different reactions with different RPA primer pairs, +: positive control. **C.** Verification of eight reactions with different components based a combined system of CRISPR-Cas12a and ssDNA FAM-BHQ1. 1-7: different reactions with different components, *rpoB*L378R and *rpoB*_wild types were added to reactions 3 and 7, respectively, PAGE: polyacrylamide gel electrophoresis.

To improve the CRISPR-Cas12a detection efficiency, we optimized fluorescence detection conditions. These included incubation time, the concentration of RPA primers and ssDNA FAM-BHQ1 fluorescence reporter as well as the ratio of crRNA and LbCas12a. After identifying the mutants using the CRISPR-Cas12a fluorescence assay, we next tested if the incubation time of reaction could be reduced. The results (Figure 3A and B) showed that at the incubation time of 15-minutes (min) there was fluorescence under UV, LED blue light, and the strongest fluorescence was observed at the incubation of 25 min. In optimizing concentrations of ssDNA FAM-BHQ1 reporters, the fluorescence could be observed under no excitation light in the reaction with 4.8 μM ssDNA FAM-BHQ1, and this concentration was used in the subsequent fluorescence assays (Figure 4A and B). In optimizing concentrations of RPA primers we found that the primers at 0.24 μM gave the best result among the tested concentrations (Figure 5A and B). In particular, the fluorescence signal could be observed remarkably by naked eyes in Figure 5A. The ratio of crRNA to LbCas12a might play a critical role in improving cleavage activity and detection sensitivity. As shown in Figure 6A and B, at the ratios of 1:2 and 2:1 similar fluorescence signal emitted in real-time fluorescence curve, but we later observed that he ratio of 2:1 of crRNAs to LbCas12a gave a better result when crRNA was at 400 nM in the reaction system.

**Figure 3.**
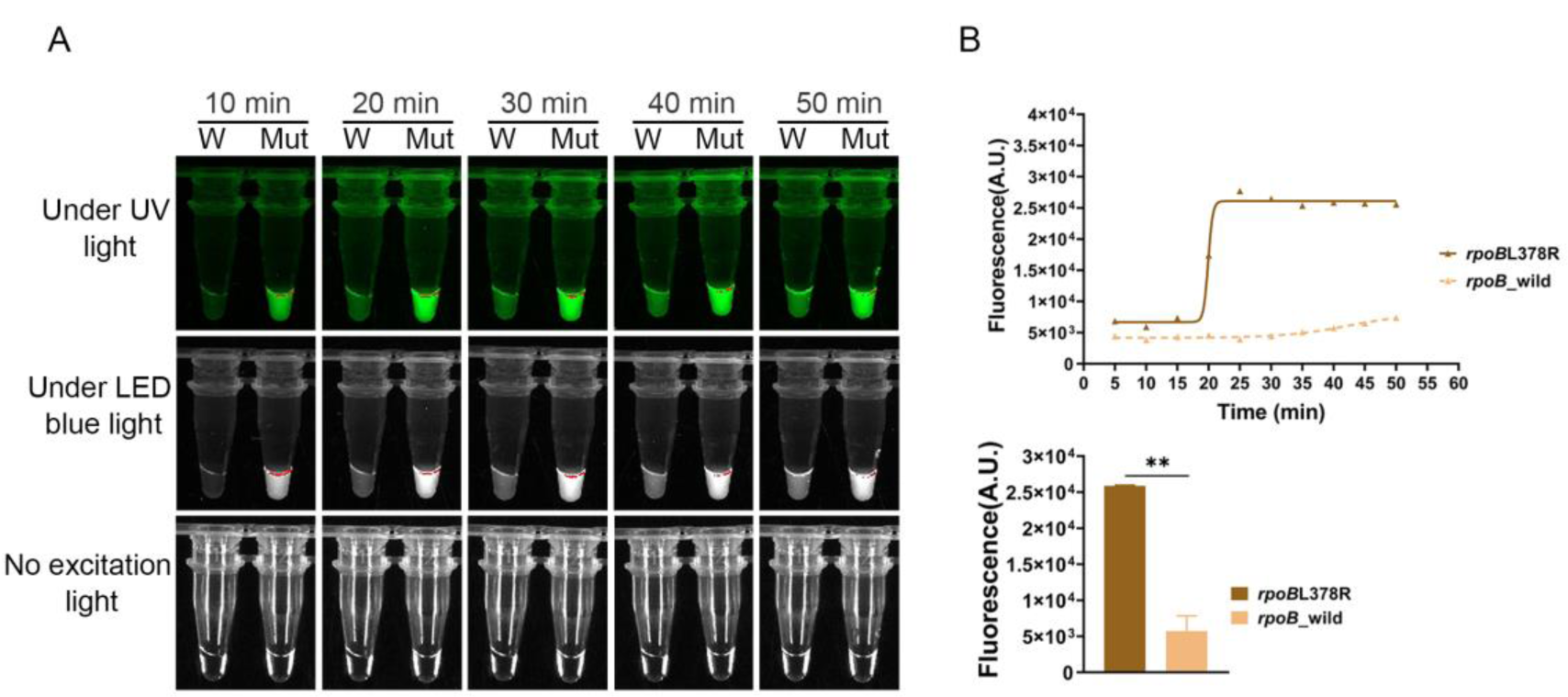
Optimization of incubation time for fluorescence detection. **A.** Determination of the best incubation time by observing fluorescence signal under UV light, LED blue light and no excitation light. W: *rpoB*_wild, Mut: *rpoB*L378R. **B.** Real-time fluorescence detection for *rpoB*_wild and *rpoB*L378R reactions with a multimode plate reader.

**Figure 4.**
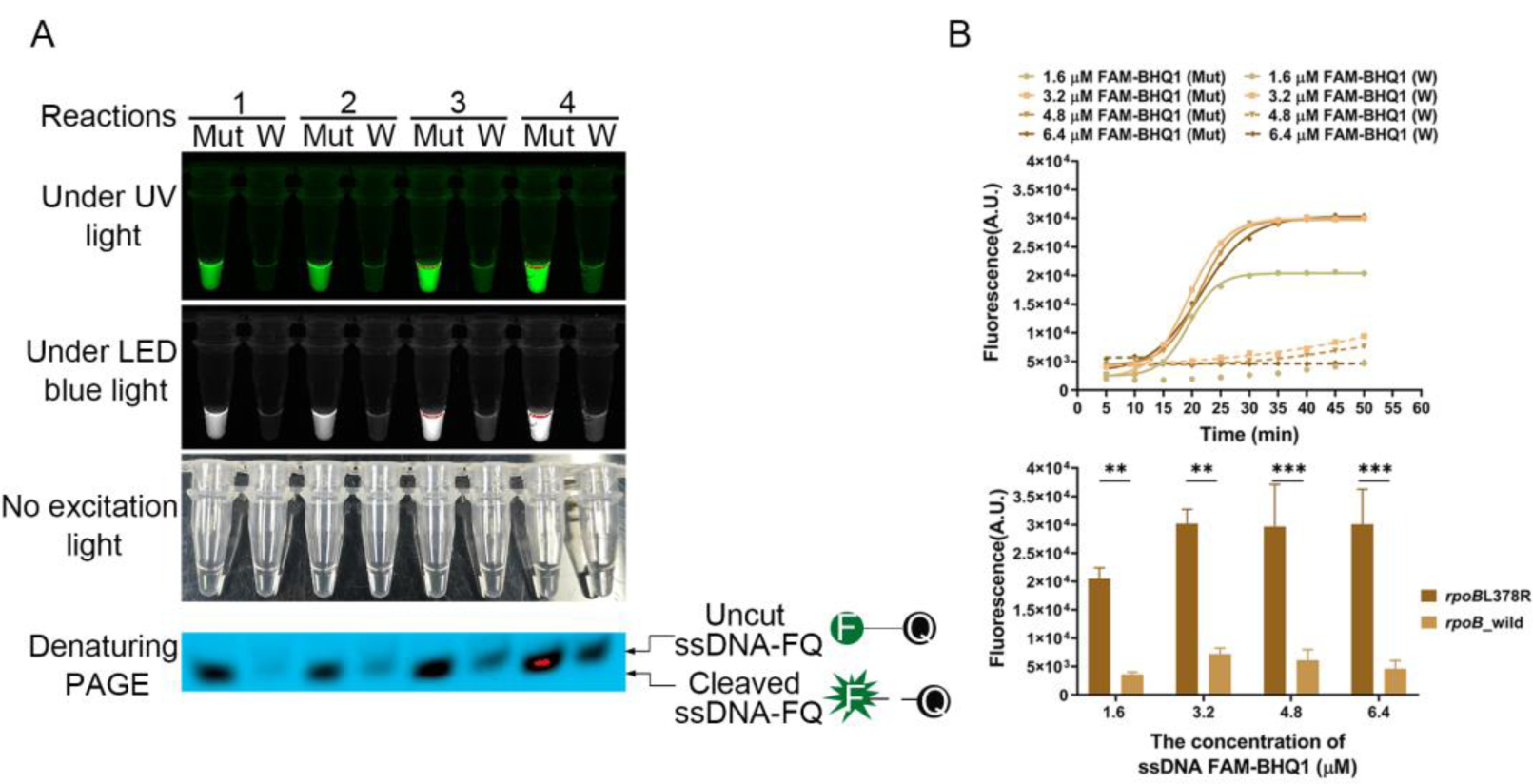
Optimization of concentrations of ssDNA FAM-BHQ1 reporter for fluorescence detection. **A.** Determination of the best concentration of ssDNA FAM-BHQ1 reporter by observing fluorescence signal under UV light, LED blue light and no excitation light. W: *rpoB*_wild, Mut: *rpoB*L378R. 1-4: different reactions with 1.6, 3.2, 4.8 and 6.4 μM ssDNA FAM-BHQ1. **B.** Real-time fluorescence detection for *rpoB*_wild and *rpoB*L378R reactions with different concentration of ssDNA FAM-BHQ1 reporter by a multimode plate reader.

**Figure 5.**
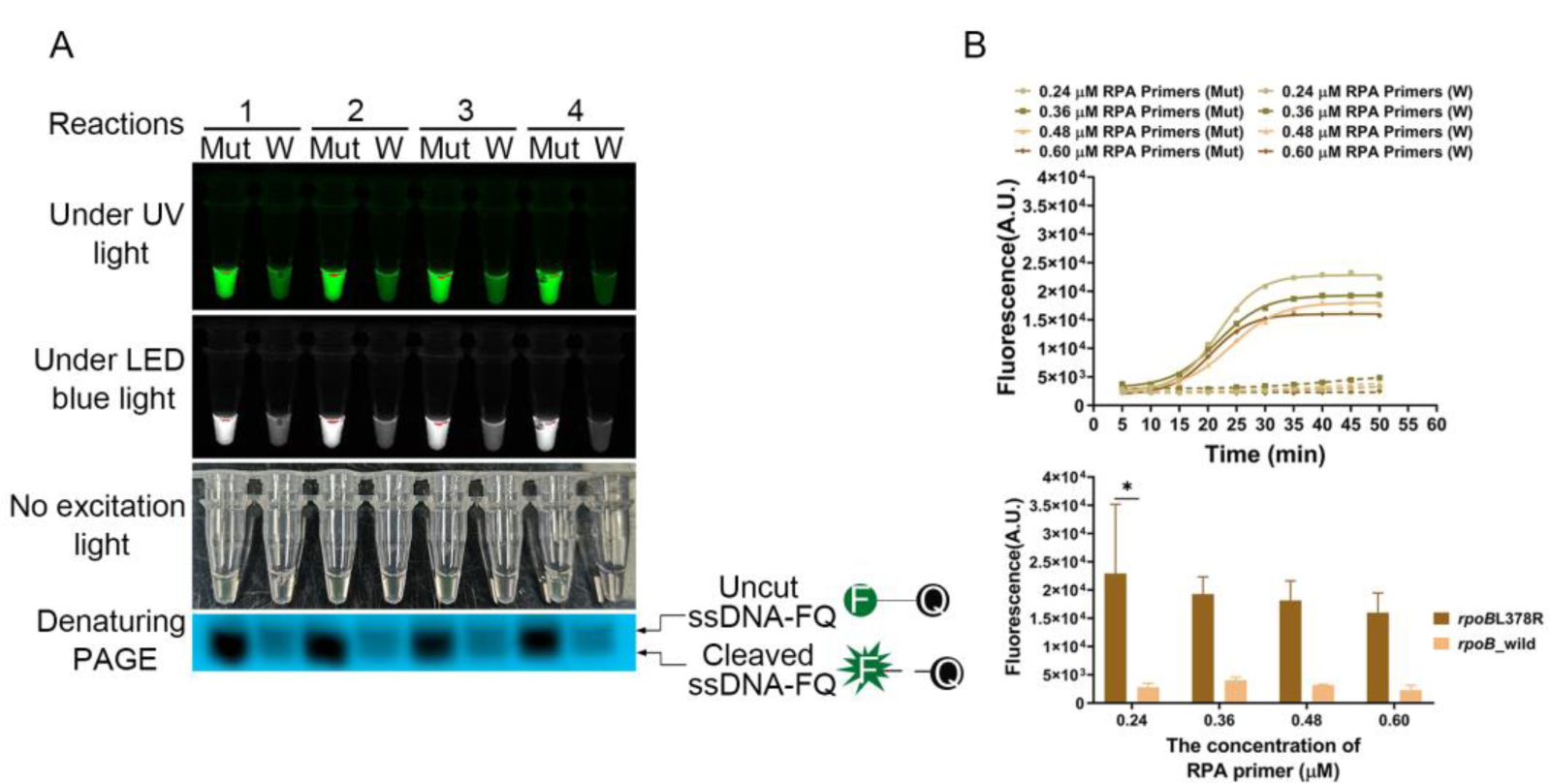
Optimization of concentrations of RPA primers for fluorescence detection. **A.** Determination of the best concentration of RPA primers by observing fluorescence signal under UV light, LED blue light and no excitation light. W: *rpoB*_wild, Mut: *rpoB*L378R. 1-4: different reactions with different molar mass of RPA primers. **B.** Real-time fluorescence detection for *rpoB*_wild and *rpoB*L378R reactions with different concentration of RPA primers by multimode plate reader.

**Figure 6.**
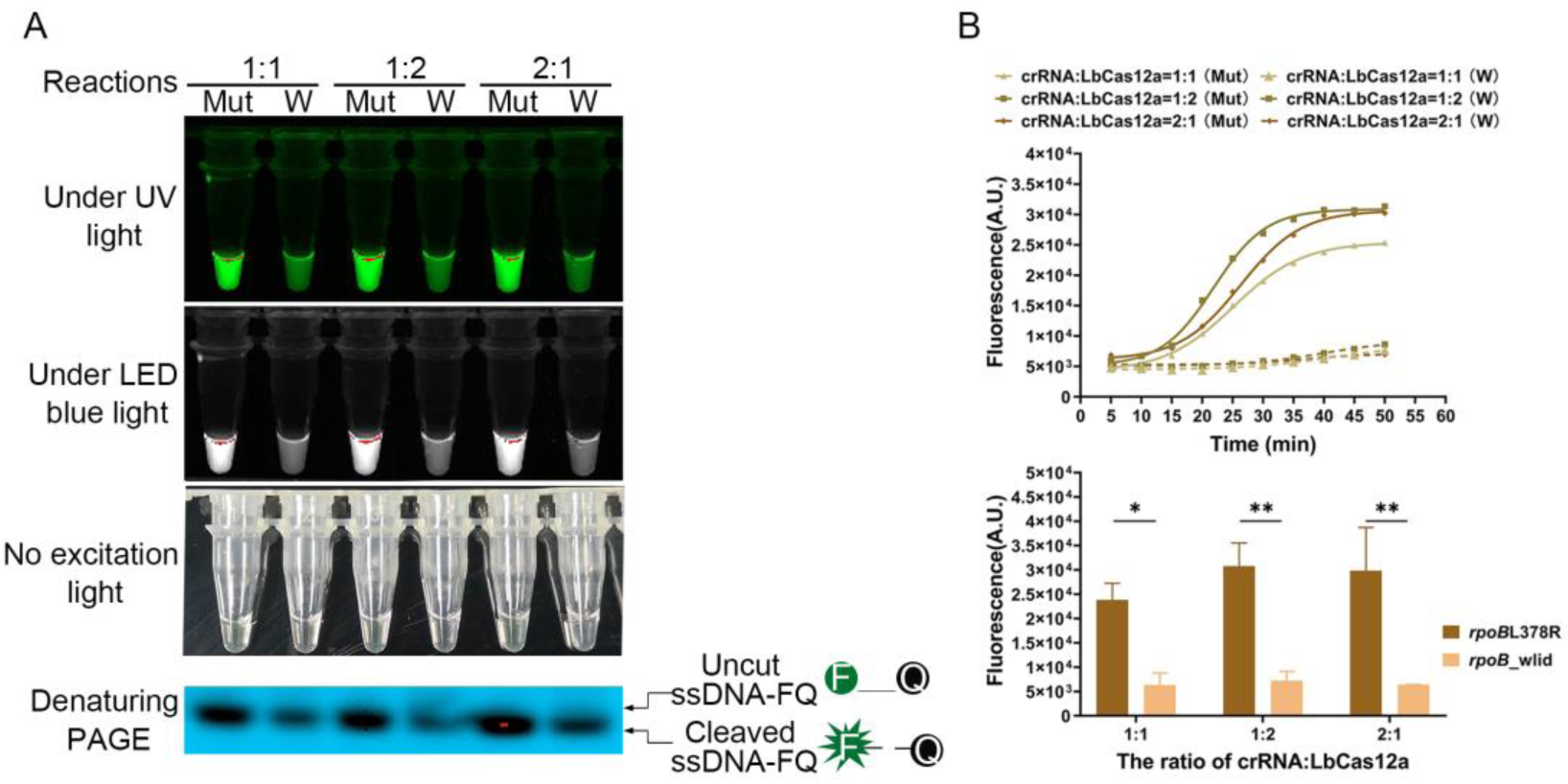
Optimization of the ratios of crRNA and LbCas12a for fluorescence detection. **A.** Determination of the ratios of crRNA and LbCas12a by fluorescence intensity under UV light, LED blue light and no excitation light. 1:1, 1:2 and 2:1 based on 400 nM of crRNA and LbCas12a. W: *rpoB*_wild, Mut: *rpoB*L378R. **B.** Real-time fluorescence detection for *rpoB*_wild and *rpoB*L378R reactions with different ratio of crRNA and LbCas12a by multimode plate reader.

### 2.3 Detection of *ropB*L378R with a CRISPR-Cas12a-original lateral flow detection assay

To reduce instrument dependence and increase efficiency, we combined lateral flow strip with CRISPR-Cas12a. In the lateral flow assay, 5’-FAM and 3’-biotin-labeled 12-nt ssDNA reporter were combined with a lateral flow strip for fluorescence intensity readout (Figure 7A) [25, 29]. Similar to the fluorescence detection assay, with lateral flow strips we quickly detected *rpoB*L378R from *rpoB*_wild type with high sensitivity without requiring for any signal detection instrument (Figure 7B). Band-intensity analysis showed that there was a higher ratio of the test band to control band resulted from the *rpoB*L378R samples than from the *rpoB*_wild (Figure 7B). These results together show that there are two methods of fluorescence signal detection and paper strip systems to detect *rpoB*L378R mutation with CRISPR-Cas12a, and these approaches are rapid, sensitive and specific.

**Figure 7.**
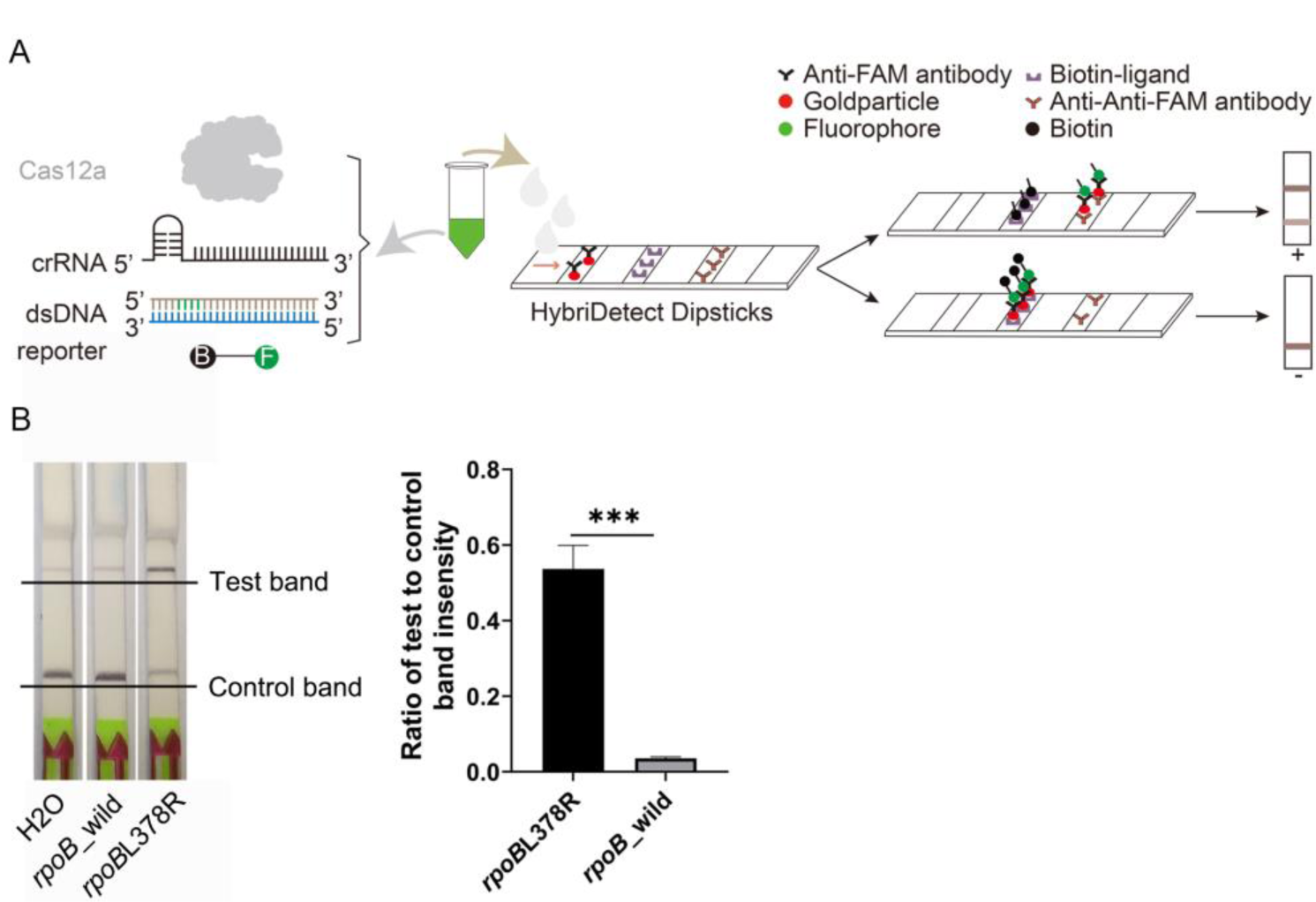
Schematic of lateral flow strip and its application for detecting *rpoB*L378R. **A.** Schematic of lateral flow strip. +: positive, -: negative. **B.** Evaluation of detection effect by lateral flow strip. Error bars represent the means ± s.d. from replicates (n=3). Two-tailed Student t-test. * p<0.05, ** p<0.01, *** p<0.001, **** p<0.0001, ns: not significant.

### 2.4 Detection of *rpoB*L378R in biological samples using CRISPR-Cas12a

We next evaluated the drug resistance mutation detection sensitivity and specificity with CRISPR-Cas12a-linked fluorescence detection and paper chromatography in different bacteria and human cells, which included HEK293T cell, *M. smegmatis*, *M. aureus*, *E. coli*, MTB *H37Rv*, and rifampicin-resistant MTB with *rpoB*L378R. Firstly, *rpoB*_wild and *rpoB*L378R fragments were mixed with *M. smegmatis*, *M. aureus* and *E. coli* genomes at molar concentration 1:1, respectively. The CRISPR-Cas12a assay showed high specificity for *rpoB*L378R (Figure 8A and B). Compared with the negative and wild-type control, the fluorescence intensity of *rpoB*L378R reaction system was the strongest, and the difference of paper chromatography was extremely significant (*P*<0.001). In detection systems of H_2_O, HEK293T, MTB *H37Rv* (*rpoB*L378R), MTB *H37Rv*, *M. smegmatis* and *M. aureus*, RPA effectively amplified *rpoB* sequence with high specificity and efficiency (Figure 8C). Fluorescence signal and paper chromatography detections based on CRISPR-Cas12a were used to identify MTB H37Rv (*rpoB*L378R). As is shown in the fluorescence detection, fluorescence intensity of *rpoB*L378R reaction system was stronger than other reaction systems (Figure 8D). Moreover, the CRISPR-Cas12a system specifically recognized *rpoB*L378R with a sensitivity of 100 aM, and there was also a significant difference in the stripe gray assay (*P* < 0.01) (Figure 8E).

**Figure 8.**
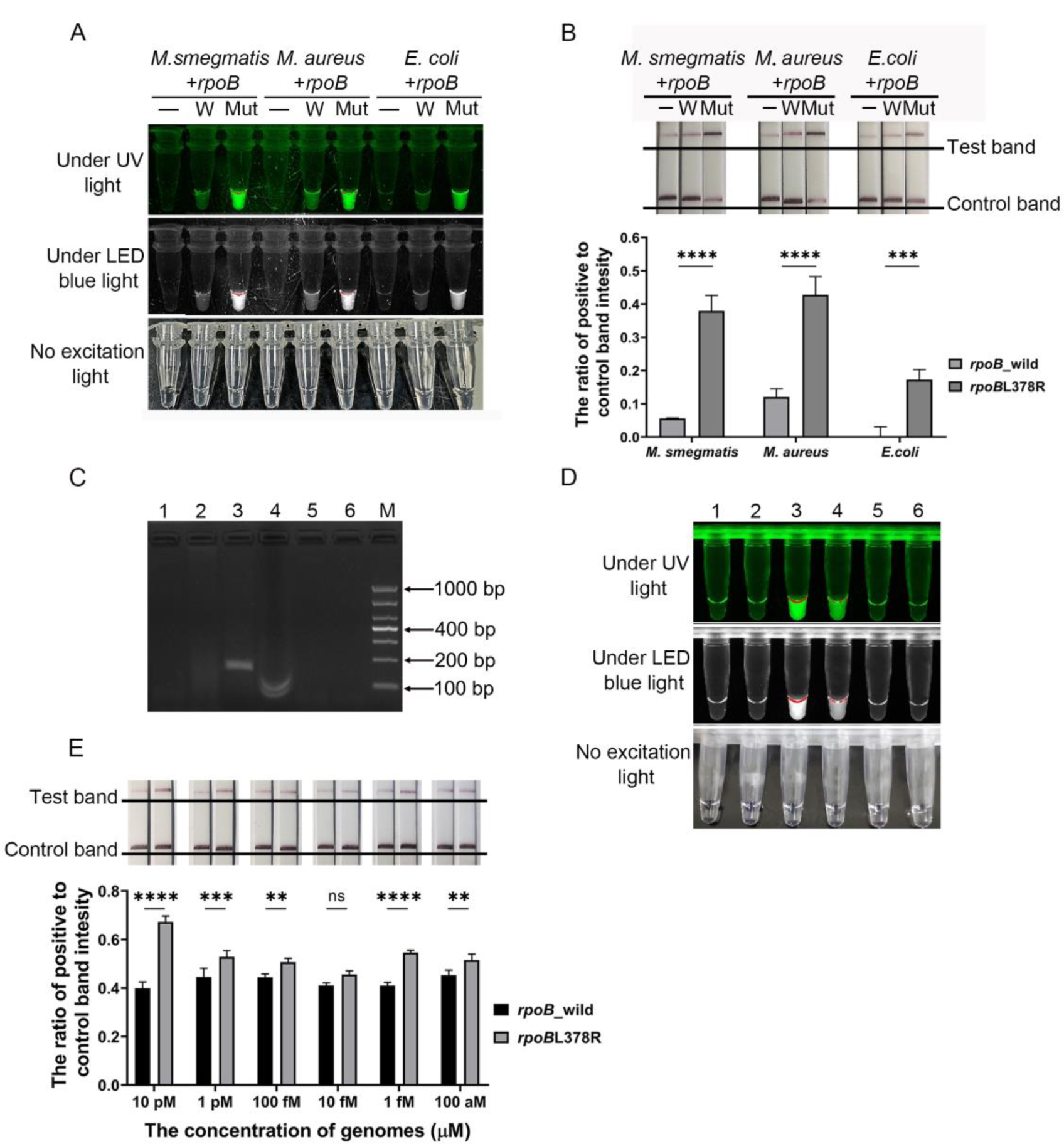
Detection of *ropB*L378R in biological samples with CRISPR-Cas12a. **A-B.** Evaluation of sensitivity and specificity in mixed system of *M. smegmatis*, *M. aureus* or *E. coli* with *rpoB*_wild or *rpoB*L378R by fluorescence signal and lateral flow strip. **C.** Evaluation of RPA amplification specificity by nucleic acid electrophoresis. 1-6: amplification reactions with H_2_O, HEK293T, MTB *H37Rv* (*rpoB*L378R), MTB *H37Rv*, *M. aureus* and *M. smegmatis*, respectively. M: DL1000 marker. **D.** Assessment of the detection specificity in MTB *H37Rv* with *rpoB*L378R. 1-6: different mixture referred to (**C**) by fluorescence intensity readout. **E.** Sensitivity evaluation of the CRISPR-Cas12a system. Error bars represent the means ±s.d. from replicates (n=3). Two-tailed Student *t*-test. * *p* < 0.05, ** *p* < 0.01, *** *p* < 0.001, **** *p*<0.0001, ns: not significant.

## 3 Discussion

In this report we show that CRISPR-Cas12a together with RPA isothermal amplification can be utilized to detect the rifampin-resistant MTB strains with the mutations such as *rpoB* L378R. This approach we developed is rapid, convenient, economical, highly specific and sensitive for identifying *rpoB*L378R; in addition, this approach does not need expensive equipment and is accurate because in this system the collateral activity of Cas12a enables highly specific and sensitive for detecting *rpoB*L378R[18, 30]. In this approach the structure of crRNA is crucial for the specificity of Cas12a-crRNA[31, 32]. It was found that cytosine residue bias, unchain temperature, purine content, and GC content at the PAM all affected the recognition efficiency and specificity of crRNA. According to previous studies, crRNAs sequences ranging from 28-35 bp were designed using CRISPR website. CrRNA design was based on the requirement that PAM sequence and drug resistance mutation point must be placed in the proximal region of PAM (1-6 bp away from PAM) to improve the detection specificity of CRISPR-Cas12a. To improve the detection specificity of CRISPR-Cas12a the mismatch base should be placed in the PAM-proximal seed region of 1-5 bp from PAM [30, 33]. By studying the recognition specificity of non-5’TTTV3’ PAM of LbCas12a, or adding additional single base mutations can also effectively improve the CRISPR-Cas12a targeting recognition specificity.

It is notable in the system that ssDNA FAM-BHQ1 fluorescent reporter cleaved by collateral activity of Cas12a is used to generate the detection results visible by producing changes in fluorescence signal of reaction system, and the signal depends on incubated time, the concentrations of the reporters and RPA primers and the ratio of crRNA to LbCas12a (Figure 3-Figure 6). We have found that 25 min incubation could maximize the detection activity of CRISPR-Cas12a system, whereas one-pot detection of RPA amplification and CRISPR-Cas12a in a single reaction system is within 60 min[12, 17]. While fluorescence reporter concentrations had a strong dose effect on the direct visual signal, the high concentrations of the reporters might produce off-target effects (Figure 4A and B).

Nonetheless, The CRISPR-Cas12a nucleic acid detection technology also has some limitations. For example, the PAM region limits its wide application, and the low concentration of fluorescent reporter also makes the fluorescence detection method unable to observe the results under natural light. It has been found that high concentration of Cas12a protein can allow more single base mismatches, and then affect the detection specificity of CRISPR-Cas detection system. A 6-base pairing of crRNA activates the “trans cleavage” activity of Cas12a, and a smaller concentration of Cas12a and crRNA increases the minimum number of matching bases recognized by CRISPR-Cas.

This system can be further improved in some aspects including integrating an ultrabright fluorescent nanocable into the detection system that has a 1000-fold lower limitation of detection[34], or optimizing the structure of crRNA, thereby reducing the amount of non-specific fluorescence reporters. In addition, since CRISPR-Cas12a-based detection only requires for controlling the temperature of incubation and RPA amplification at 37 or 39 °C, it is possible to use the minimal equipment or water bath and ubiquitous smart phone technology to record and report the detection results[12, 22, 35-37].

The reason that we choose *rpoB* gene in this study is that the target of RIF is the β-subunit of bacterial DNA-dependent RNA polymerase, which is encoded by the *rpoB* gene. At the genetic level, the majority of RIF resistance is due to the accumulation of mutations within an 81-bp region of *rpoB*, termed the rifampicin resistance determinant region (RRDR). Mutations within this region account for up to 98% of the RIF resistance observed. The strong correlation between genotypic changes in this region resulting in phenotypic resistance makes the RRDR an optimal target for the design of rapid molecular diagnostics. Recent genomic studies identified several *rpoB* non-RRDR mutations that co-occurred with RRDR mutations in clinical isolates and may confer fitness compensation. *rpoB* non-RRDR mutations could be utilized as additional molecular markers for predicting the fitness of clinical rifampin-resistant M. tuberculosis strains, and the fitness cost of rifampicin-resistance mutations was attributable to their direct influence on RNAP activity. L378R mutation is the most common non-RRDR mutation.

Notably, due to the lack of continuing horizontal gene transfer, a large majority of drug resistance phenotypes in *M. tuberculosis* are caused by chromosomal mutations. Rifampicin resistance primarily through *rpoB* mutations that results in alterations to the structure of the RIF-binding pocket and confer rifampicin resistance by decreasing the binding affinity of RIF to RNAP. In addition to resistance, *rpoB* mutations confer bacterial fitness costs by either directly decreasing the transcriptional efficacy of RNAP or indirectly altering genome-wide transcriptional profiles. RRDR mutations lead to structural or surface electrostatic potential change in the catalytic centre of RNAP which may reduce transcriptional efficiency and bacterial fitness. Notably, most of these mutations are located in loop regions. Loops are the most flexible parts of a protein, and mutations in the loop regions could contribute to modulation or diversification of protein functions. For RNAP, the rearrangement of the loop region near the active site plays an important role in its transcriptional activity.

In clinical application, it is critical in detecting the drug-resistant mutants in order to treat disease effectively. The detection system we developed presents a highly sensitive and specificity in detection of *rpoB*L378R as it did not have detectable off-target effects on other sequences (e.g., HEK293 T cell, *M. smegmatis*, *M. aureus* and *E. coli*, Figure 8C-E). This outcome may attribute to high recognition specificity of crRNA[38]. In addition, the cost of a detection reaction is estimated at ∼$0.7 and can be greatly decreased when scaled-up for bulk production in our study. Furthermore, disposable microfluidics chip or lyophilized reagents might further improve and develop the CRISPR-Cas12a-based gene detection to enable simpler and higher-efficiency point-of-care at economically underdeveloped areas or outdoors [39-41]. Moreover, this approach with the CRISPR-Cas12a system makes it possible to detect thousands or millions of nucleotide sequence, point mutations or pathogens with unlimited amounts of tested samples[23, 42].

## Conclusion

A simple, rapid, economical, highly specific, highly sensitive and visualized single-point mutations detection method we developed in this study based on CRISPR-Cas12a for MTB drug-resistant gene *rpoB*L378R establishes a solid foundation for developing a new detection kit to screen TB drug resistance gene mutations of RIF.

## Data availability statement

The datasets used and analyzed during the current study are available from the corresponding authors on reasonable request.

## Author Contributions

Yanhui Yang initiated the study. Li Yang, Yanhui Yang, Minghai Shan and Hetian Lei designed the paper. Li Yang performed all experiments with assistance from Yue Zhu, Kai Ma, Yuma Yang, Zhaoyuan Hui and Yanyan Qin. Li Yang, Xiaoyu Li and Jing Tang performed data analysis. Li Yang, Zhaoyuan Hui, Hetian Lei, Minghai Shan and Yanhui Yang wrote and revised the manuscript. All authors have read and approved the final manuscript.

---

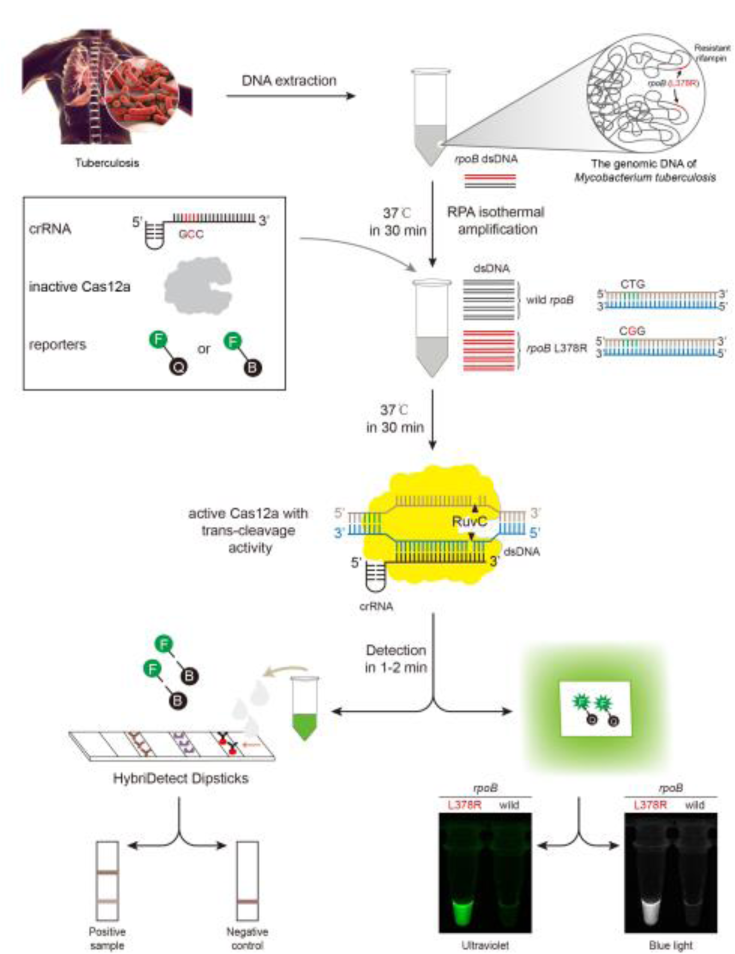
For Table of Contents Only

## Reference

1. Sotgiu, G., R. Centis, L. D’Ambrosio, and G.B. Migliori, Tuberculosis treatment and drug regimens. Cold Spring Harb Perspect Med, 2015. 5(5): p. a017822.

2. Organization, W.H., World Health Organization 2021 Geneva: World Health Organization, 2021.

3. Torres Ortiz, A., J. Coronel, J. Vidal, C. Bonilla, D. Moore, R. Gilman, F. Balloux, O. Kon, X. Didelot, and L. Grandjean, Genomic signatures of pre-resistance in Mycobacterium tuberculosis. Nature communications, 2021. 12(1): p. 7312.

4. Welekidan, L., S. Yimer, E. Skjerve, T. Dejene, H. Homberset, T. Tønjum, and O. Brynildsrud, Mycobacterium tuberculosisWhole Genome Sequencing of Drug Resistant and Drug Susceptible Isolates From Tigray Region, Ethiopia. Frontiers in microbiology, 2021. 12: p. 743198.

5. Ma, P., T. Luo, L. Ge, Z. Chen, X. Wang, R. Zhao, W. Liao, and L. Bao, Compensatory effects of M. tuberculosis rpoB mutations outside the rifampicin resistance-determining region. Emerg Microbes Infect, 2021. 10(1): p. 743–752.

6. Ramirez, M., K. Cowart, P. Campbell, G. Morlock, D. Sikes, J. Winchell, and J. Posey, Rapid detection of multidrug-resistant Mycobacterium tuberculosis by use of real-time PCR and high-resolution melt analysis. Journal of clinical microbiology, 2010. 48(11): p. 4003–9.

7. Cao, L., X. Cui, J. Hu, Z. Li, J.R. Choi, Q. Yang, M. Lin, L. Ying Hui, and F. Xu, Advances in digital polymerase chain reaction (dPCR) and its emerging biomedical applications. Biosens Bioelectron, 2017. 90: p. 459–474.

8. Radmard, S., S. Reid, P. Ciryam, A. Boubour, N. Ho, J. Zucker, D. Sayre, W.G. Greendyke, B.A. Miko, M.R. Pereira, S. Whittier, D.A. Green, and K.T. Thakur, Clinical Utilization of the FilmArray Meningitis/Encephalitis (ME) Multiplex Polymerase Chain Reaction (PCR) Assay. Front Neurol, 2019. 10: p. 281.

9. Shragai, T., S.E. Smith-Jeffcoat, M. Koh, M.C. Schechter, P.A. Rebolledo, V. Kasinathan, Y. Wang, A. Hoffman, H. Miller, A. Tejada-Strop, S. Jain, A. Tamin, J.L. Harcourt, N.J. Thornburg, P. Wong, M. Medrzycki, J.M. Folster, V. Semenova, E. Steward-Clark, J. Drobenuic, C. Biedron, R.J. Stewart, J. da Silva, H.L. Kirking, J.E. Tate, and C.C.-E.R.G.-F. Team, Epidemiologic, immunologic, and virus characteristics in patients with paired SARS-CoV-2 serology and reverse transcription polymerase chain reaction testing. J Infect Dis, 2021. 225 (2): p. 229–237

10. Habte, D., M. Melese, N. Hiruy, Z. Gashu, D. Jerene, F. Moges, S. Yifru, B. Girma, Y. Kassie, Y.K. Haile, P.G. Suarez, and B. Tessema, The additional yield of GeneXpert MTB/RIF test in the diagnosis of pulmonary tuberculosis among household contacts of smear positive TB cases. Int J Infect Dis, 2016. 49: p. 179–84.

11. Cai, Z., Z. Wang, C. Liu, D. Shi, D. Li, M. Zheng, H. Han-Zhang, A. Lizaso, J. Xiang, J. Lv, W. Wu, Z. Zhang, Z. Zhang, F. Yuan, S. He, and J. Sun, Detection of Microsatellite Instability from Circulating Tumor DNA by Targeted Deep Sequencing. J Mol Diagn, 2020. 22(7): p. 860–870.

12. Ding, X., K. Yin, Z. Li, R.V. Lalla, E. Ballesteros, M.M. Sfeir, and C. Liu, Ultrasensitive and visual detection of SARS-CoV-2 using all-in-one dual CRISPR-Cas12a assay. Nat Commun, 2020. 11(1): p. 4711.

13. Miao, F., J. Zhang, N. Li, T. Chen, L. Wang, F. Zhang, L. Mi, J. Zhang, S. Wang, Y. Wang, X. Zhou, Y. Zhang, M. Li, S. Zhang, and R. Hu, Rapid and Sensitive Recombinase Polymerase Amplification Combined With Lateral Flow Strip for Detecting African Swine Fever Virus. Front Microbiol, 2019. 10: p. 1004.

14. Vanhomwegen, J., A. Kwasiborski, A. Diop, L. Boizeau, D. Hoinard, M. Vray, R. Bercion, B. Ndiaye, A. Dublineau, S. Michiyuki, J.C. Manuguerra, V. Sauvage, D. Candotti, A. Seck, S. Laperche, and Y. Shimakawa, Development and clinical validation of loop-mediated isothermal amplification (LAMP) assay to diagnose high HBV DNA levels in resource-limited settings. Clin Microbiol Infect, 2021. 27(12): p. 1858.e9-1858.e15.

15. He, Z., Y. Su, S. Li, P. Long, P. Zhang, and Z. Chen, Development and Evaluation of Isothermal Amplification Methods for Rapid Detection of Lethal Amanita Species. Front Microbiol, 2019. 10: p. 1523.

16. Bhattacharyya, R.P., S.G. Thakku, and D.T. Hung, Harnessing CRISPR Effectors for Infectious Disease Diagnostics. ACS Infect Dis, 2018. 4(9): p. 1278–1282.

17. Ding, X., K. Yin, Z. Li, and C. Liu, All-in-One Dual CRISPR-Cas12a (AIOD-CRISPR) Assay: A Case for Rapid, Ultrasensitive and Visual Detection of Novel Coronavirus SARS-CoV-2 and HIV virus. bioRxiv, 2020. undefined(undefined): p. undefined

18. Gootenberg, J.S., O.O. Abudayyeh, M.J. Kellner, J. Joung, J.J. Collins, and F. Zhang, Multiplexed and portable nucleic acid detection platform with Cas13, Cas12a, and Csm6. Science, 2018. 360(6387): p. 439-444.

19. Aquino-Jarquin, G., CRISPR-Cas14 is now part of the artillery for gene editing and molecular diagnostic. Nanomedicine, 2019. 18: p. 428–431.

20. Huang, Y., D. Gu, H. Xue, J. Yu, Y. Tang, J. Huang, Y. Zhang, and X. Jiao, Rapid and Accurate Campylobacter jejuni Detection With CRISPR-Cas12b Based on Newly Identified Campylobacter jejuni-Specific and-Conserved Genomic Signatures. Front Microbiol, 2021. 12: p. 649010.

21. Ding, R., J. Long, M. Yuan, X. Zheng, Y. Shen, Y. Jin, H. Yang, H. Li, S. Chen, and G. Duan, CRISPR/Cas12-Based Ultra-Sensitive and Specific Point-of-Care Detection of HBV. Int J Mol Sci, 2021. 22(9): p. undefined.

22. Broughton, J.P., X. Deng, G. Yu, C.L. Fasching, V. Servellita, J. Singh, X. Miao, J.A. Streithorst, A. Granados, A. Sotomayor-Gonzalez, K. Zorn, A. Gopez, E. Hsu, W. Gu, S. Miller, C.Y. Pan, H. Guevara, D.A. Wadford, J.S. Chen, and C.Y. Chiu, CRISPR-Cas12-based detection of SARS-CoV-2. Nat Biotechnol, 2020. 38(7): p. 870–874.

23. Ackerman, C.M., C. Myhrvold, S.G. Thakku, C.A. Freije, H.C. Metsky, D.K. Yang, S.H. Ye, C.K. Boehm, T.F. Kosoko-Thoroddsen, J. Kehe, T.G. Nguyen, A. Carter, A. Kulesa, J.R. Barnes, V.G. Dugan, D.T. Hung, P.C. Blainey, and P.C. Sabeti, Massively multiplexed nucleic acid detection with Cas13. Nature, 2020. 582(7811): p. 277-282.

24. Myhrvold, C., C.A. Freije, J.S. Gootenberg, O.O. Abudayyeh, H.C. Metsky, A.F. Durbin, M.J. Kellner, A.L. Tan, L.M. Paul, L.A. Parham, K.F. Garcia, K.G. Barnes, B. Chak, A. Mondini, M.L. Nogueira, S. Isern, S.F. Michael, I. Lorenzana, N.L. Yozwiak, B.L. MacInnis, I. Bosch, L. Gehrke, F. Zhang, and P.C. Sabeti, Field-deployable viral diagnostics using CRISPR-Cas13. Science, 2018. 360(6387): p. 444-448.

25. Chen, J.S., E. Ma, L.B. Harrington, M. Da Costa, X. Tian, J.M. Palefsky, and J.A. Doudna, CRISPR-Cas12a target binding unleashes indiscriminate single-stranded DNase activity. Science, 2018. 360(6387): p. 436-439.

26. Harrington, L.B., D. Burstein, J.S. Chen, D. Paez-Espino, E. Ma, I.P. Witte, J.C. Cofsky, N.C. Kyrpides, J.F. Banfield, and J.A. Doudna, Programmed DNA destruction by miniature CRISPR-Cas14 enzymes. Science, 2018. 362(6416): p. 839-842.

27. Kellner, M.J., J.G. Koob, J.S. Gootenberg, O.O. Abudayyeh, and F. Zhang, SHERLOCK: nucleic acid detection with CRISPR nucleases. Nat Protoc, 2019. 14(10): p. 2986–3012.

28. Mustafa, M.I. and A.M. Makhawi, SHERLOCK and DETECTR: CRISPR-Cas Systems as Potential Rapid Diagnostic Tools for Emerging Infectious Diseases. J Clin Microbiol, 2021. 59(3): p. undefined.

29. Moreno-Mateos, M.A., J.P. Fernandez, R. Rouet, C.E. Vejnar, M.A. Lane, E. Mis, M.K. Khokha, J.A. Doudna, and A.J. Giraldez, CRISPR-Cpf1 mediates efficient homology-directed repair and temperature-controlled genome editing. Nat Commun, 2017. 8(1): p. 2024.

30. Kim, H., W.J. Lee, Y. Oh, S.H. Kang, J.K. Hur, H. Lee, W. Song, K.S. Lim, Y.H. Park, B.S. Song, Y.B. Jin, B.H. Jun, C. Jung, D.S. Lee, S.U. Kim, and S.H. Lee, Enhancement of target specificity of CRISPR-Cas12a by using a chimeric DNA-RNA guide. Nucleic Acids Res, 2020. 48(15): p. 8601–8616.

31. Nguyen, L.T., B.M. Smith, and P.K. Jain, Enhancement of trans-cleavage activity of Cas12a with engineered crRNA enables amplified nucleic acid detection. Nat Commun, 2020. 11(1): p. 4906.

32. Ke, Y., S. Huang, B. Ghalandari, S. Li, A.R. Warden, J. Dang, L. Kang, Y. Zhang, Y. Wang, Y. Sun, J. Wang, D. Cui, X. Zhi, and X. Ding, Hairpin-Spacer crRNA-Enhanced CRISPR/Cas13a System Promotes the Specificity of Single Nucleotide Polymorphism (SNP) Identification. Adv Sci (Weinh), 2021. 8(6): p. 2003611.

33. Swarts, D.C., J. van der Oost, and M. Jinek, Structural Basis for Guide RNA Processing and Seed-Dependent DNA Targeting by CRISPR-Cas12a. Mol Cell, 2017. 66(2): p. 221–233 e4.

34. Liu, L., Z. Wang, Y. Wang, J. Luan, J.J. Morrissey, R.R. Naik, and S. Singamaneni, Plasmonically Enhanced CRISPR/Cas13a-Based Bioassay for Amplification-Free Detection of Cancer-Associated RNA. Adv Healthc Mater, 2021. 10(20): p. e2100956.

35. Xie, S., D. Tao, Y. Fu, B. Xu, Y. Tang, L. Steinaa, J.D. Hemmink, W. Pan, X. Huang, X. Nie, C. Zhao, J. Ruan, Y. Zhang, J. Han, L. Fu, Y. Ma, X. Li, X. Liu, and S. Zhao, Rapid Visual CRISPR Assay: A Naked-Eye Colorimetric Detection Method for Nucleic Acids Based on CRISPR/Cas12a and a Convolutional Neural Network. ACS Synth Biol, 2021. 11(1): p. 383–396

36. Yin, K., V. Pandian, K. Kadimisetty, C. Ruiz, K. Cooper, J. You, and C. Liu, Synergistically enhanced colorimetric molecular detection using smart cup: a case for instrument-free HPV-associated cancer screening. Theranostics, 2019. 9(9): p. 2637–2645.

37. Chen, W., H. Yu, F. Sun, A. Ornob, R. Brisbin, A. Ganguli, V. Vemuri, P. Strzebonski, G. Cui, K. Allen, S. Desai, W. Lin, D. Nash, D. Hirschberg, I. Brooks, R. Bashir, and B. Cunningham, Mobile Platform for Multiplexed Detection and Differentiation of Disease-Specific Nucleic Acid Sequences, Using Microfluidic Loop-Mediated Isothermal Amplification and Smartphone Detection. Analytical chemistry, 2017. 89(21): p. 11219–11226.

38. Kim, H., W. Lee, Y. Oh, S. Kang, J. Hur, H. Lee, W. Song, K. Lim, Y. Park, B. Song, Y. Jin, B. Jun, C. Jung, D. Lee, S. Kim, and S. Lee, Enhancement of target specificity of CRISPR-Cas12a by using a chimeric DNA-RNA guide. Nucleic acids research, 2020. 48(15): p. 8601–8616.

39. Li, N., Y. Lu, J. Cheng, and Y. Xu, A self-contained and fully integrated fluidic cassette system for multiplex nucleic acid detection of bacteriuria. Lab on a chip, 2020. 20(2): p. 384–393.

40. Song, J., M. Mauk, B. Hackett, S. Cherry, H. Bau, and C. Liu, Instrument-Free Point-of-Care Molecular Detection of Zika Virus. Analytical chemistry, 2016. 88(14): p. 7289–94.

41. Song, J., C. Liu, M. Mauk, J. Peng, T. Schoenfeld, and H. Bau, A Multifunctional Reactor with Dry-Stored Reagents for Enzymatic Amplification of Nucleic Acids. Analytical chemistry, 2018. 90(2): p. 1209–1216.

42. Vervoort, Y., A.G. Linares, M. Roncoroni, C. Liu, J. Steensels, and K.J. Verstrepen, High-throughput system-wide engineering and screening for microbial biotechnology. Curr Opin Biotechnol, 2017. 46: p. 120–125.

